# Early transcriptome analysis of the brown streak virus–cassava pathosystem provides molecular insights into virus susceptibility and resistance

**DOI:** 10.1101/100552

**Authors:** Ravi B. Anjanappa, Devang Mehta, Michal J. Okoniewski, Alicja Szabelska, Wilhelm Gruissem, Hervé Vanderschuren

**Author notes:** These authors contributed equally to this work (Ravi B. Anjanappa, Devang Mehta). Author for correspondence: / /.

## Abstract

Cassava brown streak viruses (CBSVs) are responsible for significant cassava yield losses in eastern sub–Saharan Africa. In the present work, we inoculated CBSV–susceptible and –resistant cassava varieties with a mixed infection of CBSVs using top-cleft grafting. Virus titres in grafted scions were monitored in a time course experiment in both varieties. We performed RNA-seq of the two cassava varieties at the earliest time-point of full infection in the susceptible scions. Genes encoding proteins in RNA silencing and salicylic acid pathways were regulated in the susceptible cassava variety but transcriptional changes were limited in the resistant variety. After infection, genes related to callose deposition at plasmodesmata were regulated and callose deposition was significantly reduced in the susceptible cassava variety. We also show that β–1,3–glucanase enzymatic activity is differentially regulated in the susceptible and resistant varieties. The differences in transcriptional responses to CBSV infection indicate that resistance involves callose deposition at plasmodesmata but does not trigger typical anti-viral defence responses. A meta-analysis of the current RNA-seq dataset and selected, previously reported, host–potyvirus and virus-cassava RNA-seq datasets revealed comparable host responses across pathosystems only at similar time points after infection or infection of a common host.

**HIGHLIGHT:** Our results suggest that resistance to CBSV in cassava involves callose deposition at the plasmodesmata and our meta-analysis of multiple virus-crop RNA-seq studies suggests that conserved responses across different host-virus systems are limited and depend greatly on time after infection.

## INTRODUCTION

Cassava brown streak disease (CBSD) is one of the most damaging cassava virus diseases in Africa causing harvest losses of up to 70% in susceptible varieties (Hillocks et al., 2001; Legg et al., 2011). Ongoing CBSD outbreaks are threatening cassava production in sub–Saharan Africa (Legg et al., 2011; Legg et al., 2014) and the recent spread of CBSD into central Africa (Bigirimana et al., 2011; Mulimbi et al., 2012) indicates that it could soon become a major constraint to cassava production in all sub–Saharan cassava growing regions.

The characteristic symptoms of CBSD include leaf chlorosis and dry hard necrosis in roots, thus affecting both the quality and yield of the edible storage roots. CBSD is caused by two phylogenetically distinct cassava brown streak virus species: *Cassava brown streak virus* (CBSV) and *Ugandan cassava brown streak virus* (UCBSVs) (Mbanzibwa et al., 2011), collectively referred to as CBSVs. CBSVs are (+) ssRNA viruses belonging to the genus *Ipomovirus*, family Potyviridae (Monger et al., 2001; Mbanzibwa et al., 2009), and are transmitted by whiteflies (Maruthi et al., 2005). They consist of a ~9 kb genome with a single open reading frame encoding a large poly-protein, which is co- and post–translationally cleaved into ten viral proteins (Winter et al., 2010). CBSV and UCBSV differ in virulence in both controlled and field conditions, with CBSV generally accumulating to higher titres in most host genotypes (Winter et al., 2010; Mohammed et al., 2012; Kaweesi et al., 2014; Ogwok et al., 2015). Two recent field analyses of different cassava genotypes for the presence of CBSV and UCBSV found that both viruses co–occurred in a majority of tested genotypes (Ogwok et al., 2015; Kaweesi et al., 2014).

The release of a cassava reference genome (Prochnik et al., 2012) has opened up new opportunities to perform large–scale characterization of the cassava transcriptome and proteome (Allie et al., 2014; Maruthi et al., 2014; Vanderschuren et al., 2014). Cassava varieties differing in resistance against cassava geminiviruses (CMGs) were recently analysed in a time course experiment to identify pathways that are differentially regulated during South African cassava mosaic virus (SACMV) infection (Allie et al., 2014). Similarly, a late (one year post–infection) time point comparative transcriptome analysis was reported of transcripts differentially expressed in a susceptible variety (Albert) and a variety (Kaleso) classified as moderately resistant but displaying mild CBSD symptoms after CBSV infection (Maruthi et al., 2014). The late time point study found no regulation in cassava homologs of known R-gene analogues, any NBS-LRR genes, nor genes encoding proteins of the anti-viral RNA silencing pathway. We recently characterised KBH 2006/18 and KBH 2006/26, which are two resistant elite breeding lines with immunity to a mixed infection of CBSV and UCBSV (Anjanappa et al., 2016). Interestingly, KBH 2006/18 and KBH 2006/26 show no symptoms or hypersensitive response upon infection, even at 16-weeks post graft-inoculation (Anjanappa et al., 2016).

Here we used two cassava varieties contrasting for CBSV resistance (i.e. 60444 and KBH2006/18) to characterize the transcriptome response at an early time-point of CBSV and UCBSV co-infection. We report an in-depth characterisation of pathways and genes regulated in compatible and incompatible cassava–ipomovirus interactions and present biochemical evidence to explain the inhibition of virus intercellular movement in the resistant breeding line KBH 2006/18. Our data provide new insights into the cassava-CBSV interaction and identify commonalities in plant responses to virus infection.

## METHODS

### Plant material and virus inoculation

The elite CBSV-resistant cassava breeding variety KBH 2006/18 was obtained from IITA (Tanzania) and the susceptible variety 60444 from the ETH cassava germplasm collection. Cassava plants were grown under greenhouse conditions (27°C, 16h light, 60% humidity). Infected 60444 stem cuttings with mixed infection of the CBSV isolate TAZ-DES-01 (KF878104) and the UCBSV isolate TAZ-DES-02 (KF878103) were used to inoculate KBH 2006/18 and 60444 plants using a previously described top-cleft grafting method (Moreno et al., 2011; Anjanappa et al., 2016). KBH 2006/18 and 60444 plants were also top-cleft grafted onto virus-free 60444 rootstocks as mock controls. Individual leaves (the second leaf from the point of graft union) were collected from scions of CBSV- and mock-infected plants at 28 days after grafting (dag) and at 60 dag (Fig. S1).

### RT-qPCR

RNA was extracted from cassava leaves using a modified CTAB RNA extraction protocol (Moreno et al., 2011). One μg of DNaseI-treated total RNA was reverse-transcribed using the RevertAid First Strand cDNA Synthesis Kit (Thermo-Fisher) according to the manufacturer’s instructions. RT-qPCR was performed with SYBR Green dye (Thermo-Fisher) using a 7500 Fast Real Time PCR System (Applied Biosystems). Three independent biological replicates were analysed for each time-point. The relative expression level of individual genes in each sample was calculated as previously described (Moreno et al., 2011; Vanderschuren et al., 2012). Primers used to validate differentially expressed genes and to quantitate virus titres are listed in Table S1.

### Illumina RNA-Seq and data analysis

Library preparation and sequencing were performed as follows: 5 μg of total RNA extract was treated with DNase according to manufacturer instructions (Qiagen, RNase-free DNase Kit) and subsequently column purified using the RNeasy Plant Mini Kit (Qiagen). RNA quality was assessed using a Qubit^®^ (1.0) Fluorometer (Thermo-Fisher) as well as a Bioanalyzer 2100 (Agilent). Samples with a RNA integrity number (RIN) above 6 were used for sequencing. One μg of total RNA was polyA enriched and mRNA libraries were synthesized using a TruSeq RNA Sample Prep Kit v2 (Illumina). Cluster generation was done using 10 pM of pooled normalized libraries on the cBOT with TruSeq PE Cluster Kit v3–cBot–HS (Illumina) and subsequently Illumina HiSeq 2000 sequencing was performed to generate the reads.

### Sequence assembly, mapping to the cassava genome and identification of differentially expressed genes (DEGs)

The raw read files were trimmed using the trim sequence tool in the CLC Genomics Workbench v8.5 and settings declared in Notes S1. Unique trimmed reads were mapped to the *M. esculenta* reference genome AM560-2 v6.0 available on Phytozome (www.phytozome.jgi.doe.gov) using the transcriptomics toolbox in the CLC Genomics Workbench v8.5. Empirical analysis of differentially expressed reads was performed using default parameters to identify differentially expressed genes (DEGs) between control non-infected and CBSVs-infected scions. DEGs were then filtered based on a FDR-corrected P-value of <0.01 and a fold–change of ≥2. DEGs were annotated in the CLC Workbench using the Phytozome GFF3 annotation file (www.phytozome.jgi.doe.gov).

### Identification of de novo assembled transcripts (dnATs)

Read mapping of trimmed reads was performed using the transcriptomics toolbox in the CLC Genomics Workbench v8.5 with settings described in Notes S1 but with “Maximum number of hits for a read = 10”, allowing for more stringent detection of unmapped reads. Unmapped reads for each cassava variety were pooled and assembled using Trinity (Grabherr et al., 2011) with default settings. Assembled contigs were then size-filtered and contigs greater than 1000 nt were classified as *de novo* assembled transcripts (*dnATs*). Differential expression of *dnATs* between infected and mock-infected samples was determined using STAR (Dobin et al., 2013).

### Determination of full-length virus genomes from RNA-Seq analysis

Unmapped reads (described previously) were assembled into contigs using the CLC Genomics Workbench *de novo* assembly tool. The assembled contigs were BLASTed against published CBSV (HG965221.1) and UCBSV (HG965222.1) genomes to identify full length virus genomes. Two full length genomes corresponding to CBSV (TAZ-DES-01; KF878104) and UCBSV (TAZ-DES-02; KF878103) were obtained as single assembled contigs. The two full-length viral genomes were used to map reads from each individual sample (Fig. S1) using CLC Genomics Workbench read mapping to contigs tool in order to estimate virus read counts per sample and read count distribution across the virus genomes.

### β–1,3–glucanase enzymatic assay

Leaf samples from three independent cassava scions were collected and soluble proteins were extracted from 100 mg of leaf powder in 300 μl of extraction buffer (20 mM HEPES, pH 8.0, 5 mM MgCl2 and 1 tablet/50 ml buffer of EDTA–free protease inhibitor) and quantitated using the BCA assay. The amount of glucose content released by the activity of β-1,3-glucanase in a known amount of soluble proteins in presence of the substrate laminarin (Laminaria digitata, Sigma–Aldrich) was then measured spectrophotometrically at 540 nm (Ramada et al., 2010). β-1,3-glucanase specific activity was subsequently calculated, with 1 unit (U) of activity representing the amount of enzyme required to produce 1 μmol of reducing sugar per minute.

### Quantification of plasmodesmata–associated callose

Leaf sections from uninfected and infected scions were stained overnight with Aniline blue (Merck) using a previously described method (Guenoune–Gelbart et al., 2008). Stained leaf samples were then examined using a Zeiss LSM 780 laser confocal microscope with a 40X water–immersion objective. Callose accumulation was detected at an excitation of 405 nm and emission between 413 and 563nm was recorded. Callose quantification and deposit counting was performed as described by Zavaliev & Epel (2015) for eight replicate images per plant variety and treatment condition.

### Pfam-based meta-analysis

For Pfam-based meta-analysis, RNA-Seq DEG data were downloaded from published reports (Allie et al., 2014; Rubio et al., 2015 a, b; Maruthi et al., 2014) and filtered using an absolute fold-change of ≥2. Pfam annotations for genomes of each species were obtained from Phytozome annotation files (www.phytozome.jgi.doe.gov). Commonly regulated Pfams were determined by constructing a 5-set Venn diagram and the resulting Pfam overlaps between pairs of sets were used to calculate a pairwise similarity matrix.

## RESULTS

### CBSV inoculation induces greater transcriptome modulation in susceptible cassava

Following successful grafting of 60444 and KBH2006/18 virus–free scions onto rootstocks with a mixed infection of CBSV (TAZ–DES–01) and UCBSV (TAZ–DES–02) (CBSV-infection in short), we performed a time course sampling of the scions at 16, 22 and 28 days after grafting (dag) (Fig. S1). We assessed virus titres in inoculated and mock scions at the selected time points. Virus infection reached homogenous titre levels across the three biological replicates only at 28 dag in inoculated 60444 scions (Fig. S2). No virus was detected in KBH 2006/18 scions (Fig. S2), consistent with previous results (Anjanappa et al., 2016). Notably, we did not observe CBSD symptoms in any inoculated plant at 28 dag. Samples from 60444 and KBH2006/18 scions at 28 dag were selected for transcriptome analysis.

Gene expression in 60444 and KBH 2006/18 from non-inoculated and CBSV-inoculated scions collected at 28 dag was analysed using Illumina HiSeq 2000 sequencing. Total RNA from the second emergent leaf immediately after the graft-union was used for RNA-Seq analysis, with three independent replicates per condition and plant variety. A total of 1,254,619,280 reads with 100-bp paired-ends were generated from the 12 samples, ranging from 77 to 121 million reads per sample. On average, 93.4% of the total reads were successfully mapped to the cassava reference genome (*M. esculenta* AM560-2 v6.0). The reads not mapped to the reference genome were assembled de novo using Trinity (Grabherr et al., 2011). The size-selected Trinity-assembled contigs, referred to as *de novo* assembled transcripts (*dn*ATs), were subsequently blasted against the NCBI nucleotide collection to provide gene descriptions. The *dn*ATs were then used for differential gene expression analysis using STAR (Dobin et al., 2013). None of the *dn*ATs appeared to be differentially regulated in compatible and incompatible cassava-CBSV pathosystems (Table S2).

A total of 1,292 genes in 60444 and 585 genes in KBH 2006/18 were differentially expressed at 28 dag, reflecting a higher number of modulated genes in the compatible host-virus interaction (Table S3). Over 68% of all differentially expressed genes (DEGs) were up-regulated in variety 60444 while only 30% were up-regulated in KBH 2006/18. A total of 158 DEGs were common between both varieties, and a major proportion (99 DEGs) of these were down-regulated (Fig. 1a). Among the DEGs found in both plant varieties, only 12 genes were differently regulated (i.e. up-regulated in 60444 and down-regulated in KBH 2006/18 or vice-versa) (Fig.1a).

**Fig. 1:**
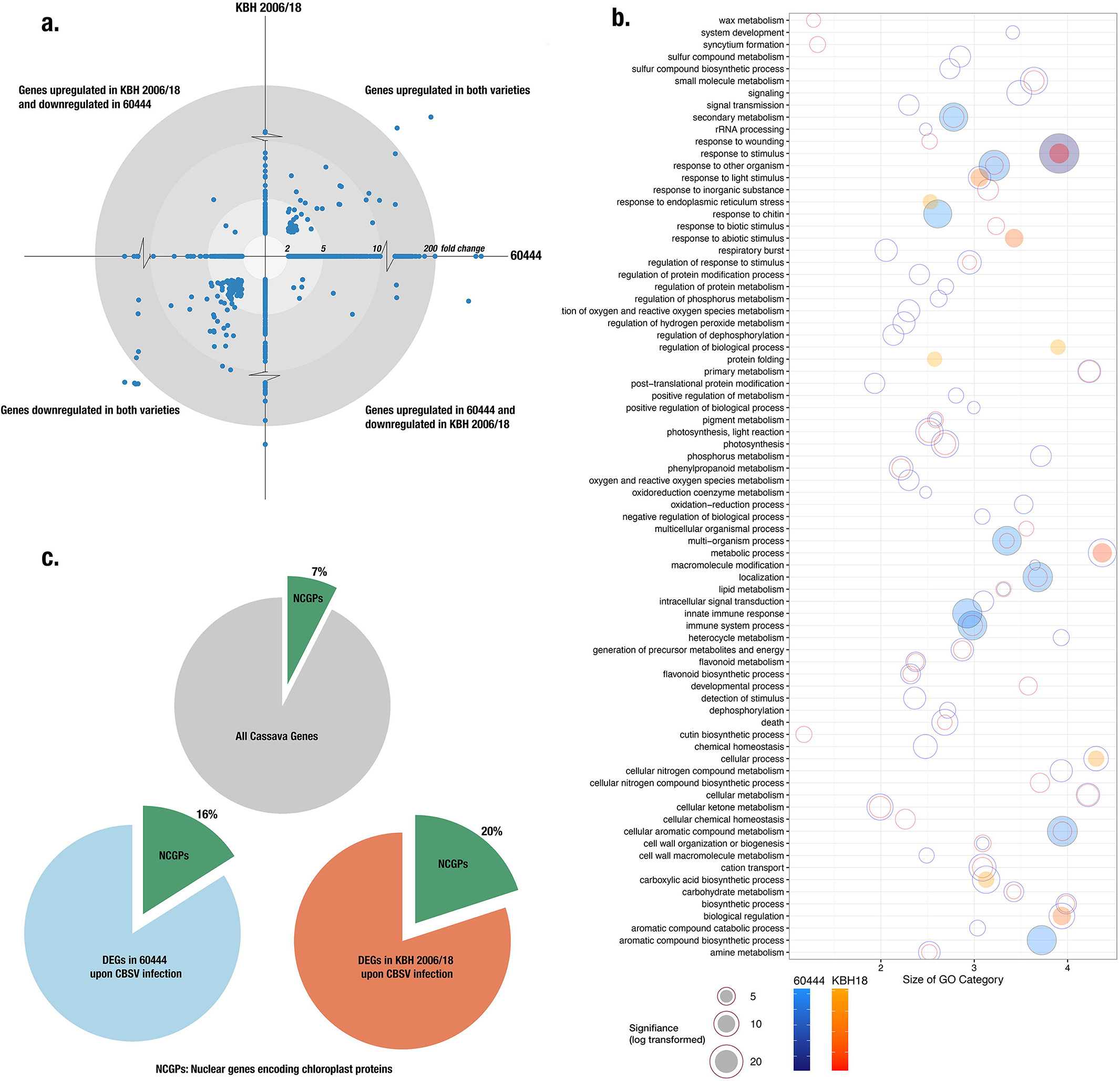
Gene regulation in susceptible (60444) and resistant (KBH 2006/18) cassava varieties at 28 dag and mixed CBSV and UCBSV infection. **a.** Fold change of differentially expressed genes (DEGs) (FDR<0.05, fold change ≥2) in both varieties. Fold change of DEGs in 60444 form the X–coordinate and in KBH 2006/18 form the Y–coordinate. **b.** Functional characterization of DEGs based on gene ontology (GO) of biological processes. Significantly enriched GO categories in 60444 and KBH 2006/18 were first reduced based on semantic similarity (REVIGO; revigo.irb.hr) and then plotted. Blue circles represent GO categories enriched in 60444 and orange circles those in KBH 2006/18; overlapping blue and orange circles represent GO categories enriched in both 60444 and KBH 2006/18. Circles for the top 10 significantly enriched categories are shaded. **c.** Proportion of nuclear-encoded chloroplast genes (NECGs) differentially regulated after infection. 6% of the cassava genome consists of NECGs while 16% of the 60444 and 20% of the KBH 2006/18 DEGs after infection are NECGs.

### Comparison and validation of RNA-Seq data

The RNA-Seq expression results of three significantly regulated genes (*PLASMODESMATA LOCATED PROTEIN 1(PDLP1), CALRETICULIN _1B_ (CRT_1B_), HEAT SHOCK PROTEIN 17.6 (HSP17.6)*) and one non-significantly regulated gene (*CALRETICULIN _3_ (CRT_3_)*), were confirmed using RT-qPCR on the same RNA samples used for sequencing. Fold-change values (comparing infected and non-infected samples) for these RNAs were similar to those observed in the RNA-Seq analysis (Fig. S3). We used unmapped reads to *de novo* assemble the full-length virus genomes of CBSV (TAZ–DES–01) and UCBSV (TAZ–DES–02). Full-length virus genome contigs were used for read counts to estimate virus abundance in each sample used for RNA-Seq (Fig. S4). Read counts detected for CBSV (TAZ–DES–01) and UCBSV (TAZ–DES–02) indicated that CBSV (TAZ–DES–01) was more prevalent than UCBSV (TAZ–DES–02) in two of the three inoculated 60444 scions. The read count analysis confirmed the absence of both CBSVs in the inoculated KBH2006/18 scions (Table S4).

### Gene sets enriched in resistant and susceptible cassava

We performed Gene Set Enrichment Analysis using the PlantGSEA toolkit (Yi et al., 2013)) to identify significantly enriched gene ontology (GO) categories and KEGG pathways (Table S4). Significantly enriched GO categories (Biological Process) were then summarized and visualized based on semantic similarity using REVIGO (Supek et al., 2011) (Fig. 1b). As expected, genes related to stimulus response, immune system process and photosynthesis were significantly regulated in both the susceptible and resistant plant varieties (Fig. 1b).

We also examined KEGG-annotated pathways enriched after virus infection in both varieties by mapping DEGs onto enriched KEGG pathways using Pathview (Luo and Brouwer, 2013) (Table S4). In both resistant and susceptible varieties, genes in photosystem I and II as well as the light harvesting complex are down–regulated. Pathways related to phenylalanine metabolism, phenylpropanoid biosynthesis and downstream products of phenylpropanoid biosynthesis were enriched in the infected 60444 scions along with several genes involved in plant-pathogen interactions. In KBH 2006/18, the cyanoamino metabolism, starch and sucrose metabolism pathways were enriched. Interestingly, the phenylpropanoid pathway was also overrepresented in KBH 2006/18, but it showed an opposite response as compared to the infected 60444 variety.

A recent transcriptome study identified the downregulation of nuclear genes encoding chloroplast proteins (NGCPs) as a key response to PAMP (specifically, bacterial pathogen) perception (Shi et al., 2015). Considering the overrepresentation of chloroplast-related genes (TAIR GO:0009507) in our data sets, we further investigated the relative abundance of NGCP transcripts regulated by CBSV infection. In 60444, 7% of all cassava NGCPs (193 out of 2682 cassava NGCPs), which represent 15% of the total DEGs, were regulated upon infection. In KBH 2006/18, 19% of all DEGs are NGCPs (Fig. 1c). In both varieties, a majority of NGCPs were down-regulated (Table S4), indicating that CBSV inoculation induces a chloroplastic response, as observed with other PAMPs in Arabidopsis (Göhre et al., 2012; Zheng et al., 2012; de Torres Zabala et al., 2015).

### CBSV infection modulates callose deposition at plasmodesmata

In a previous study investigating the cassava-CBSV pathosystem we found that KBH 2006/18 allows transmission of CBSVs through the stem vasculature while no virus can be detected in KBH 2006/18 leaves (Anjanappa et al., 2016). This suggests that the resistance of KBH 2006/18 to CBSVs involves at least restriction of inter-cellular virus movement from vascular tissues to mesophyll cells. Here we find that transcript levels of genes involved in virus movement and callose deposition at plasmodesmata were regulated in infected 60444 but not KBH 2006/18 leaves. The transcript level of *β–1,3–GLUCANASE (BG3)* encoding an enzyme which degrades callose at the plasmodesmata (Iglesias and Meins Jr, 2000), was 9.7-fold up-regulated in 60444 (Fig. 2a). ANKYRIN REPEAT FAMILY PROTEINS (ANKs) allow viral movement through plasmodesmata by promoting callose degradation (Ueki et al., 2010). We also found an upregulation of two different cassava *ANKs*, Manes.11G035500 (3.8-fold) in 60444 and Manes.05G056700 (2.03-fold) in KBH 2006/18. The expression of *PLASMODEMATA–LOCATED PROTEIN _1_ (PDLP_1_)*, which is a positive regulator of virus cell-to-cell movement, was upregulated in infected 60444. PDLP1 binds to viral movement proteins and helps the formation of viral movement tubules at the plasmodesmata (Amari et al., 2010), thereby facilitating virus cell-to-cell movement. In contrast, the expression of a possible negative regulator of plasmodesmata permeability, *GLUCAN SYNTHASE LIKE _4_ (GSL_4_)* (Maeda et al., 2014), was also up–regulated by 2.6–fold in 60444 (Fig. 2a).

**Fig. 2:**
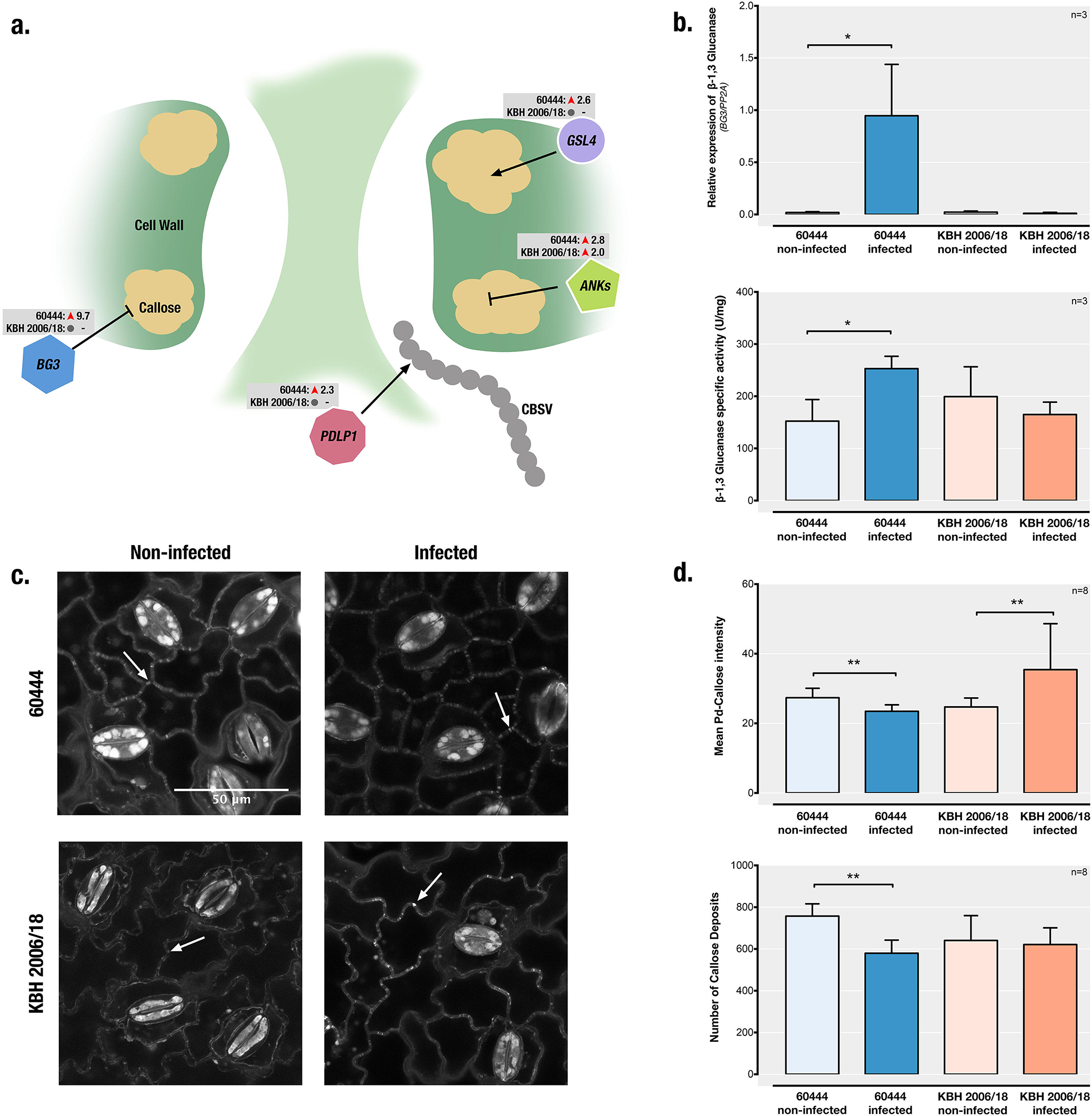
Regulation of callose deposition at plasmodesmata in susceptible (60444) and resistant (KBH 2006/18) cassava varieties after CBSV-infection. **a.** Schematic depiction of a plasmodesmata used by viruses for intercellular movement. Also shown are genes encoding key enzymes significantly regulated after infection (BG3: GLUCANASE, encoding a callose-degrading β-1,3-glucanase; GSL4: GLUCAN SYNTHASE-LIKE 4, encoding a callose synthesis enzyme; PDLP1: PLASMODEMATA-LOCATED PROTEIN 1, encoding a protein involved in virus movement across the plasmodesmata and in callose synthesis; ANKs: ANKYRIN REPEAT FAMILY PROTEINS, encoding proteins that are negative regulators of callose deposition) with their fold-changes based on RNA-Seq analysis at 28 dag. **b.** (upper panel) RT-qPCR analysis of BG3 mRNA at 60 dag. 60444 leaves were symptomatic for virus infection at this time-point. (lower panel) BG3 enzymatic activity also significantly increases after infection in 60444 and is unchanged in KBH 2006/18 at 60 dag. **c.** Aniline blue staining for detection of callose deposits. Leaf samples from 60444 were stained and visualized using a confocal laser scanning microscope at 400X magnification. Callose deposits are visible as small bright dots along the cell walls (white arrowheads). **d.** (upper panel) Quantitation of callose deposits at plasmodesmata in 60444 and KBH 2006/18 at 60 dag, observed after aniline blue staining and microscopy. Infection results in a decrease in the amount of callose deposited at the plasmodesmata in 60444 and an increase in KBH 2006/18. (lower panel) The number of callose deposits found in microscopic images. Infection also decreases in the number of callose deposits in 60444. (*, P < 0.05; **, P < 0.01, Un-paired t-test.)

We further characterized BG3 expression levels at 60 dag to follow the regulation of this gene in infected and symptomatic 60444 leaves. Up-regulation of BG3 expression was attenuated at 60 dag as the CBSV infection progressed. Consistently, no significant change of BG3 expression was observed in KBH 2006/18 leaves at 60 dag (Fig. 2b). The higher BG3 transcript levels resulted in a 38% increase in BG3 enzymatic activity in infected 60444 leaves (Fig. 2b).

In order to understand the functional implications of the simultaneous upregulation of positive (*PDLP_1_*, *GSL_4_*) and negative (*BG_3_*, *ANKs*) regulators of callose deposition upon CBSV infection, we stained callose in leaf tissue from 60444 and KBH 2006/18 at 60 days after grafting (Fig. 2c). This revealed a significant decrease in plasmodesmata-associated callose deposition in epidermal cells of infected 60444 leaves and a strong increase in the amount of plasmodesmata–associated callose deposition in KBH 2006/18.

### Salicylic acid defence and lignin biosynthesis pathways are regulated in response to CBSVs

Overall, we found that in the susceptible cassava variety 60444 defence pathways were more extensively regulated during CBSV infection as compared to the resistant variety KBH 2006/18. Transcript levels for several genes encoding the enzymes involved in lignin and SA biosynthesis were up-regulated in the compatible cassava-CBSV interaction and not regulated or down–regulated in the incompatible interaction (Fig. 3). Both SA and lignin biosynthetic pathways follow the phenylpropanoid pathway, which was enriched in our initial KEGG pathway gene–set analysis (Table S4).

**Fig. 3:**
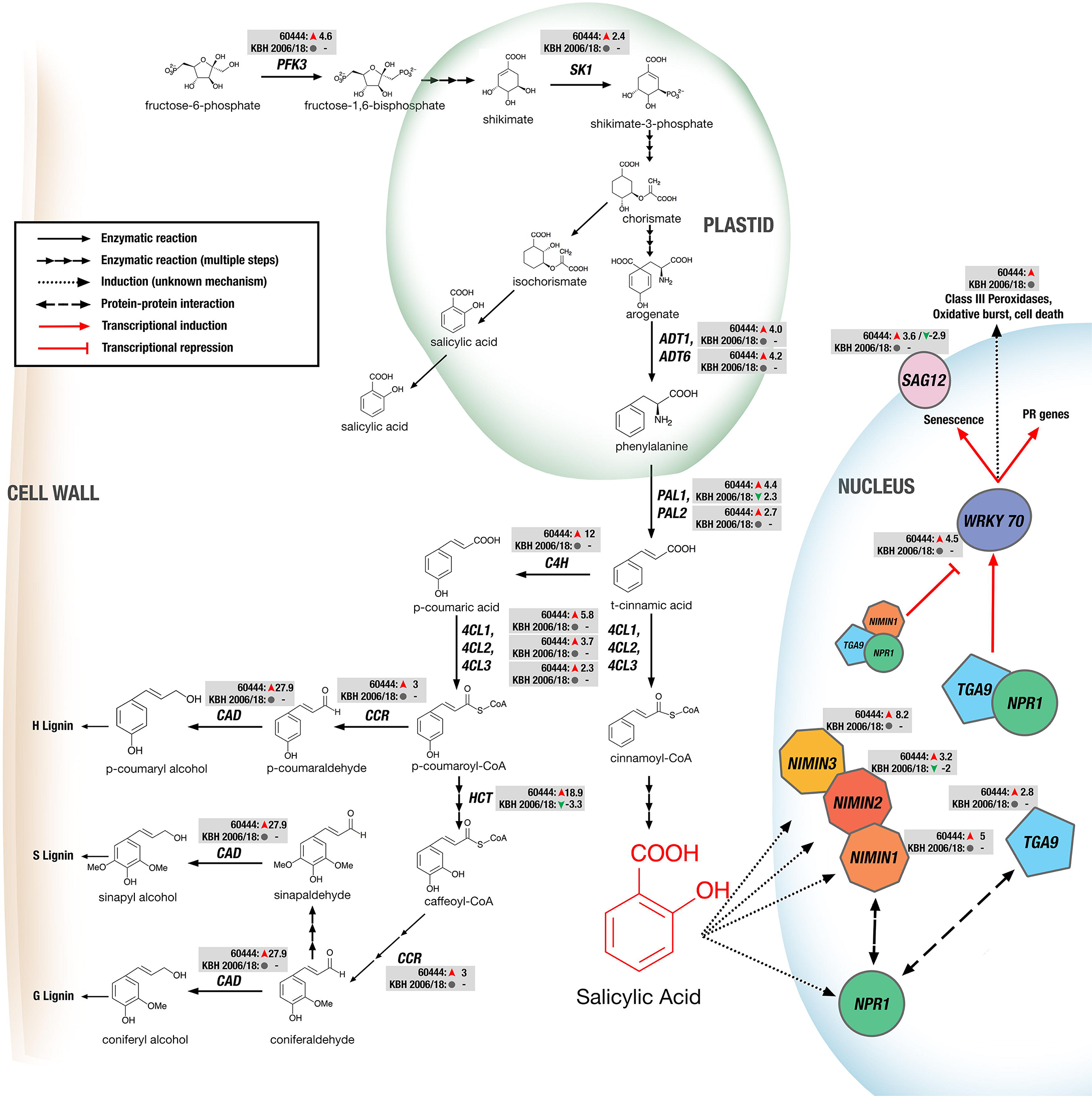
Regulation of genes encoding proteins of the salicylic acid synthesis and signalling and the lignin synthesis pathways in cassava after virus infection. Significantly regulated genes are depicted together with their mRNA fold-change based on RNA-Seq analysis in susceptible (60444) and resistant (KBH 2006/18) cassava varieties after infection. *PFK_3_*: *PHOSPHOFRUCTOKINASE _3_*; *SK_1_*: *SHIKIMATEKINASE_1_*; *ADT_1_*, *ADT_6_*: *AROGENATEDEHYDRATASE 1 & 6*; *PAL_1_*, *PAL_2_*: *PHENYLALANINE AMMONIA–LYASE 1 & 2*; *C_4_H*: *CINNAMATE–4–HYDROXYLASE*; *_4_CL_1_*, *_4_CL_2_*, *_4_CL_3_*: *4–COUMARATE–CoA LIGASE 1, 2 & 3*; *HCT*: *HYDROXYCINNAMOYL–COA SHIKIMATE/QUINATE HYDROXYCINNAMOYL TRANSFERASE*; *CCR*: *CINNAMOYL COA REDUCTASE _1_*; *CAD*: *CINNAMOYL ALCOHOL DEHYDROGENASE*; *NPR_1_*: *NON–EXPRESSOR OF PR GENES _1_*; *NIMIN_1_*, *NIMIN_2_*, *NIMIN_3_*: *NIM1–INTERACTING*; *TGA9*: *TGACG MOTIF–BINDING PROTEIN 9*; *WRKY70*: *WRKY DNA–BINDING PROTEIN 70*; *SAG_12_*: *SENESCENCE–ASSOCIATED GENE 12*.

SA biosynthesis in plants begins in the chloroplast with the shikimate pathway for synthesis of aromatic amino acids, which in turn is fed by fructose–1,6–bisphosphate (Fig. 3). Although fructose-1,6-bisphosphate is probably not a limiting precursor for SA biosynthesis, we did observe a 4.6-fold up-regulation of the PHOSPHOFRUCTOKINASE 3, a gene involved in the conversion of fructose-6-phosphate to fructose-1,6-bisphosphate in the susceptible variety. We also observed a 2.4-fold up-regulation of the *SHIKIMATE KINASE _3_*, responsible for the conversion of shikimate to shikimate–3– phosphate. Previous genetic studies in Arabidopsis and *Nicotiana benthamiana* (reviewed by Chen et al., 2009) have indicated that pathogen-induced SA synthesis is catalysed by ISOCHORISMATE SYNTHASE (ICS) to convert chorismate to isochorismate, which is then used to produce SA by an unknown mechanism. An alternative pathway for SA biosynthesis from chorismate involves its conversion to arogenate, which is the substrate of AROGENATE DEHYDRATASE (ADT) to produce phenylalanine. ADT1 and ADT6 were upregulated in 60444 after CBSV infection, suggesting that virus-induced SA synthesis in cassava, unlike in *N. benthamiana* (Catinot et al., 2008), might proceed through the phenylalanine pathway rather than via isochorismate. PHENYLALANINE AMMONIA-LYASE (PAL), which catalyses the conversion of phenylalanine to cinnamic acid, is a key enzyme in the synthesis of both lignin and SA. Indeed, quadruple *pal* mutants with only 10% PAL activity produced about 50% less SA and were more susceptible to biotic stress (Huang et al., 2010). We find that *PAL_1_* (Manes.10G047500) and *PAL_2_* (Manes.07G098700) were 4.4-fold and 2.7-fold up-regulated, respectively, in 60444. In contrast, *PAL_1_* was downregulated 2.3–fold in KBH 2006/18. Cinnamic acid is the substrate at the branch point of the SA and lignin biosynthesis pathways. Ligation of cinnamic acid to Coenzyme A by 4-COUMARATE: COA LIGASE (4CL) leads to SA biosynthesis via both peroxisomal β-oxidative and cytosolic non-oxidative pathways. The conversion of cinnamic acid first to p-coumaric acid by CINNAMATE–4–HYDROXYLASE (C4H) and then ligation of Coenzyme A to p-coumaric acid by _4_CL produces p-coumaroyl–CoA, which is a phenylpropanoid pathway precursor (Widhalm and Dudareva, 2015). In our study, *_4_CL1* (Manes.14G151400, Manes.04G095300), *_4_CL2* (Manes.11G071800), *_4_CL3* (Manes.09G127000.1) and *C_4_H* (Manes.02G227200) were all up-regulated in CBSV-infected 60444 (Fig. 3).

SA is a key regulator of plant defence responses and has been implicated in various mechanisms resulting in the hypersensitive response (HR) and systemic acquired resistance (SAR) (reviewed in Vlot et al., 2009). One such mechanism is the NPR1-dependent response (Vlot et al., 2009), which involves the SA-induced activation and nuclear localisation of NPR1 to activate the transcription of genes for pathogenesis-related proteins, including BG3. NPR1 interacts with various members of the TGA family of transcription factors, which together activate the expression of WKRY transcription factors (Vlot et al., 2009). We find that *TGA_9_* (Manes.04G004100) expression was up-regulated by 2.8-fold in CBSV-infected 60444, together with the upregulation of two cassava *WRKY70* genes (Manes.07G142400 by 4.5-fold and Manes.10G002200 by 2.5-fold). In Arabidopsis, WRKY70 is an SA-induced transcription factor that activates the expression of several pathogenesis-related genes, including genes encoding a peroxidase and senescence-associated proteins (*SAG_12_*, *SAG_21_*, *SAG_24_*, *SAG_29_*)(Li et al., 2004; Ülker et al., 2007). In CBSV-infected 60444, *SAG_12_* (Manes.16G038400) was upregulated 3.6-fold (Fig. 3, Table S3). Similarly, the genes encoding 10 out of 14 Class III peroxidases (Prxs, or PR-9 subfamily), which are involved in SA-induced pathogen defence responses (Almagro et al., 2009), were upregulated (Fig. 3).

Lignins are polymers of aromatic compounds and integral components of the plant cell wall. They have a role in defence against microbial and fungal pathogens (Vanholme et al., 2010; Miedes et al., 2014). Lignin synthesis is often induced in plant immune responses (Malinovsky et al., 2014). We find several genes encoding enzymes involved in the production of phenylpropanoid intermediates and their conversion to the three monolignols that were up–regulated after CBSV infection in 60444 (Fig. 3, Table S3).

### The antiviral RNA silencing pathway is up-regulated in the compatible cassava-CBSVs interaction

RNA silencing is a major defence mechanism against plant viruses (Wang et al., 2012), especially against RNA viruses, whose genomes and their replicative forms can be directly targeted by RNA-induced Silencing Complexes (RISCs) and DICER–LIKE (DCL) proteins (Waterhouse and Fusaro, 2006). RNA silencing is thought to begin with the synthesis and formation of long dsRNA molecules, which are substrates of DCL proteins in plants (Fig. 4) (Pumplin and Voinnet, 2013). In Arabidopsis, four DCL proteins are involved in RNA silencing, of which DCL2 and DCL4 have overlapping functions in antiviral defence (Garcia–Ruiz et al., 2010). However, DCL4 alone is sufficient for antiviral RNA silencing (Garcia-Ruiz et al., 2010). DCL2 appears to be required for the production of secondary small interfering RNAs (siRNAs) (Parent et al., 2015). We find that the expression of three cassava DCL2 genes was increased in CBSV-infected 60444 (Manes.12G002700 by 2.4-fold, Manes.12G002800 by 3.3-fold, Manes.12G003000 by 3.3-fold).

**Fig. 4:**
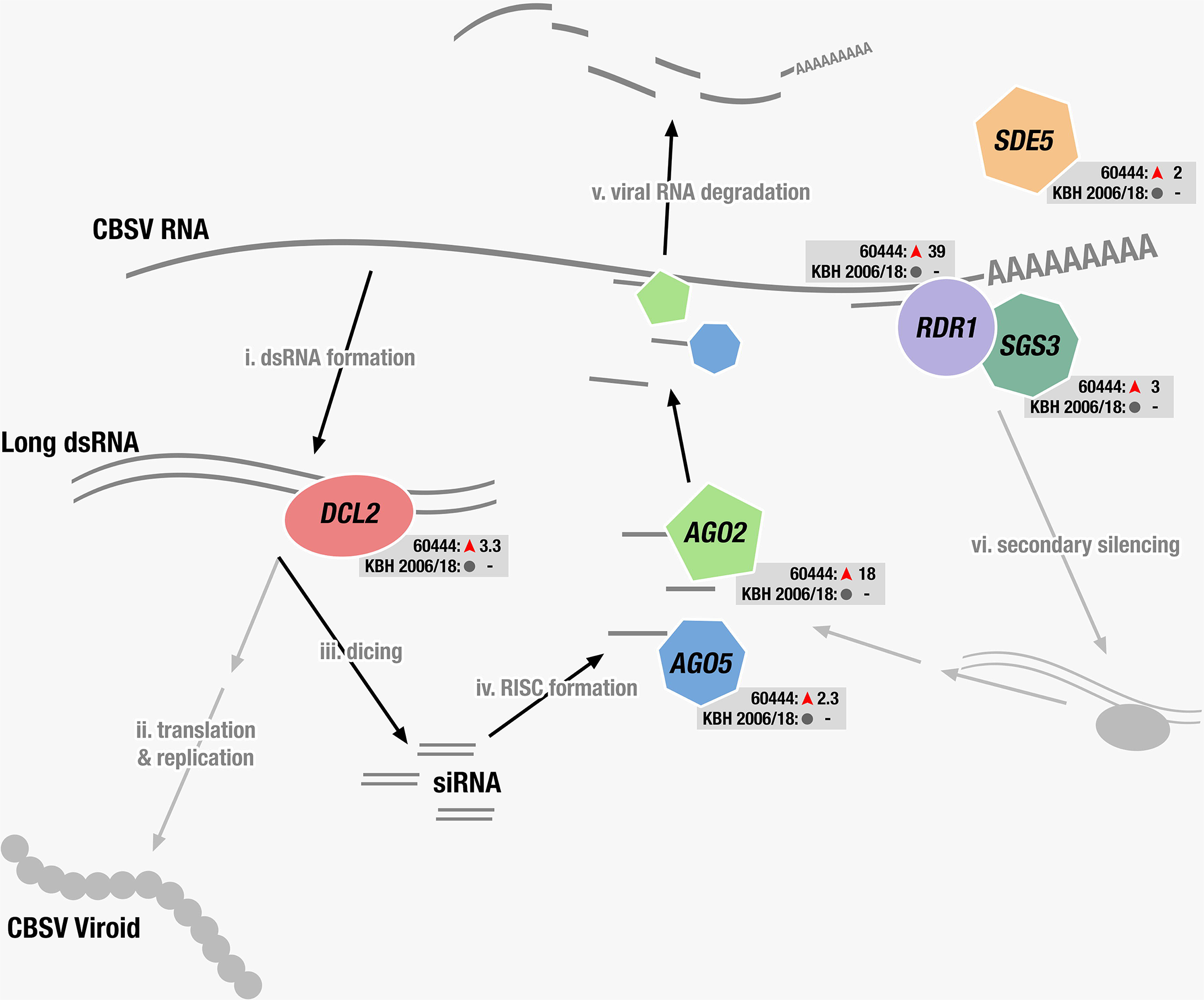
Schematic depiction of the antiviral RNA silencing pathway with DEGs encoding proteins that function in antiviral defence. Genes significantly regulated in variety 60444 after inoculation are shown together with their mRNA fold-change based on RNA-Seq analysis. These genes were not regulated in variety KBH 18/2006. DCL2: DICER–LIKE 2, AGO5: ARGONAUTE 5, AGO2: ARGONAUTE 2, RDR1: RNA DEPENDENT RNA POLYMERASE 1, SGS3: SUPRESSOR OF GENE SILENCING 3, SDE5: SILENCING DEFECTIVE 5.

ARGONAUTE (AGO) are essential proteins of the RNA Induced Silencing Complexes (RISCs), which are involved in the sequence-specific silencing of genes by RNA degradation, inhibition of translation and epigenetic modifications. Of the 10 AGO proteins known in plants, AGO2 has a broad and likely specific role in antiviral silencing (Jaubert et al., 2011; Carbonell et al., 2012; Scholthof et al., 2011). Expression of the cassava AGO2 gene (Manes.07G023100) was 17.5-fold up-regulated in CBSV-infected 60444 scions. A gene expressing AGO5 (Manes.04G011400), which is another but less well characterised AGO protein, was also up–regulated 2.3-fold. The virus-induced increased expression of *AGO_2_* and *AGO_5_* in cassava is similar to *N. benthamiana*, in which Potato Virus X infection increases *AGO_5_* expression and AGO5 acts synergistically with AGO2 to suppress virus infection to a greater degree than the other AGO proteins (Brosseau and Moffett, 2015).

RNA-DEPENDENT RNA POLYMERASE 1 (RDR1) is an important component of the post-transcriptional gene silencing (PTGS) immune response to virus infections and is involved in the defence of various classes of viruses, including potyviruses. (Rakhshandehroo et al., 2009; Garcia– Ruiz et al., 2010; Wang et al., 2010; Lee et al., 2016). RDRs catalyse the synthesis of the antisense strand in the conversion of viral RNA to a long dsRNA template that can be processed by DCLs. PTGS is often initiated by siRNAs binding to a target and recruiting RDRs, which then results in the production of further long dsRNA templates for the production of secondary siRNAs. This mechanism, also known as transitivity or silencing amplification, is largely facilitated by RDR6, and possibly also RDR1 (Wang et al., 2010). We find that expression of RDR1 (Manes.17G084800) was highly up-regulated (38.5-fold) in CBSV-infected 60444 scions (Fig. 4). The steep up-regulation of RDR1 was sustained at all three early time points (Fig. S5). The precise role of RDR1 in virus resistance has not yet been established, and one study in particular (in *N. benthamiana*) demonstrated a repressor function of RDR1 in antiviral silencing (Ying et al., 2010). It is also possible that RDR1 contributes to virus resistance by targeting host genes. Cao et al. (2014) identified 369 genes as potential RDR1 targets and validated the RDR1-dependent reduction in transcription of *PHOTOSYSTEM II LIGHT HARVESTING COMPLEX B_1.3_ (LHCB_1.3_)* gene in response to potyvirus infection. We consistently observed a 3.4-fold downregulation of *LHCB_1.3_* (Manes.17G066700) in CBSV-infected 60444 scions. However, of the 26 putative RDR1 target genes (out of the 369 genes identified by Cao et al., (2014)) regulated in our study, only 13 were down-regulated (Table S6).

The expression of the PTGS-related genes *SILENCING DEFECTIVE _5_ (SDE_5_)* and *SUPPRESSOR OF GENE SILENCING _3_ (SGS_3_)* were also significantly up-regulated in CBSV-infected 60444 (Fig. 3, Table S3) but not changed in CBSV-infected KBH 2006/18. AGO2 and RDR1, which are both up-regulated in CBSV-infected 60444, are also induced by SA signalling pathways (Lewsey et al., 2010; Jovel et al., 2011; Hunter et al., 2013; Lee et al., 2016), which is consistent with the upregulation of several cassava genes involved in SA synthesis and signalling pathways after CBSV infection.

### Similar transcript regulation in selected host-potyvirus and cassava-virus pathosystems

We performed a comparative analysis of the DEGs we detected above with other RNA-Seq studies of host-virus pathosystems to understand similarities and differences in gene expression regulation. The studies we selected involved naturally occurring crop-virus interactions with detectable host responses to potyvirus infections and comparable RNA–Seq data sets (late-stage cassava-CBSV infection, Maruthi et al., 2014; early-stage peach and late-stage apricot infection with Plum Pox Virus, Rubio et al., 2015a, b). We also included an RNA-Seq study of cassava infected with South African Cassava Mosaic Virus, a ssDNA geminivirus pathosystem (Allie et al., 2014), to compare the response of cassava to two different virus types.

We first filtered DEGs from all datasets according to our differential expression criteria (fold change >2; FDR <0.01) and used Pfam identifiers (Finn et al., 2015) matching each DEG as a common term to compare gene expression between cassava and the two Prunus species. This allowed us to construct a similarity matrix based on a pairwise similarity percentage score, where 100% denotes complete similarity (Fig. 5a, grey boxes). The greatest pairwise similarity (25.54%) was found between peach and cassava RNA-Seq datasets obtained within two-months post-infection. Interestingly, we found more common Pfam identifiers between our early time-point cassava-CBSV dataset and the early time-point cassava–SACMV dataset rather than between the two cassava-CBSV datasets. Also, the two cassava-CBSV datasets and the two Prunus datasets had a lower degree of similarity than most other pairs, indicating a strong temporal component to the modulation of host gene expression after virus infection. Our meta-analysis did not reveal a strong dependence of host gene expression changes on either RNA or DNA virus or host genus.

**Fig. 5:**
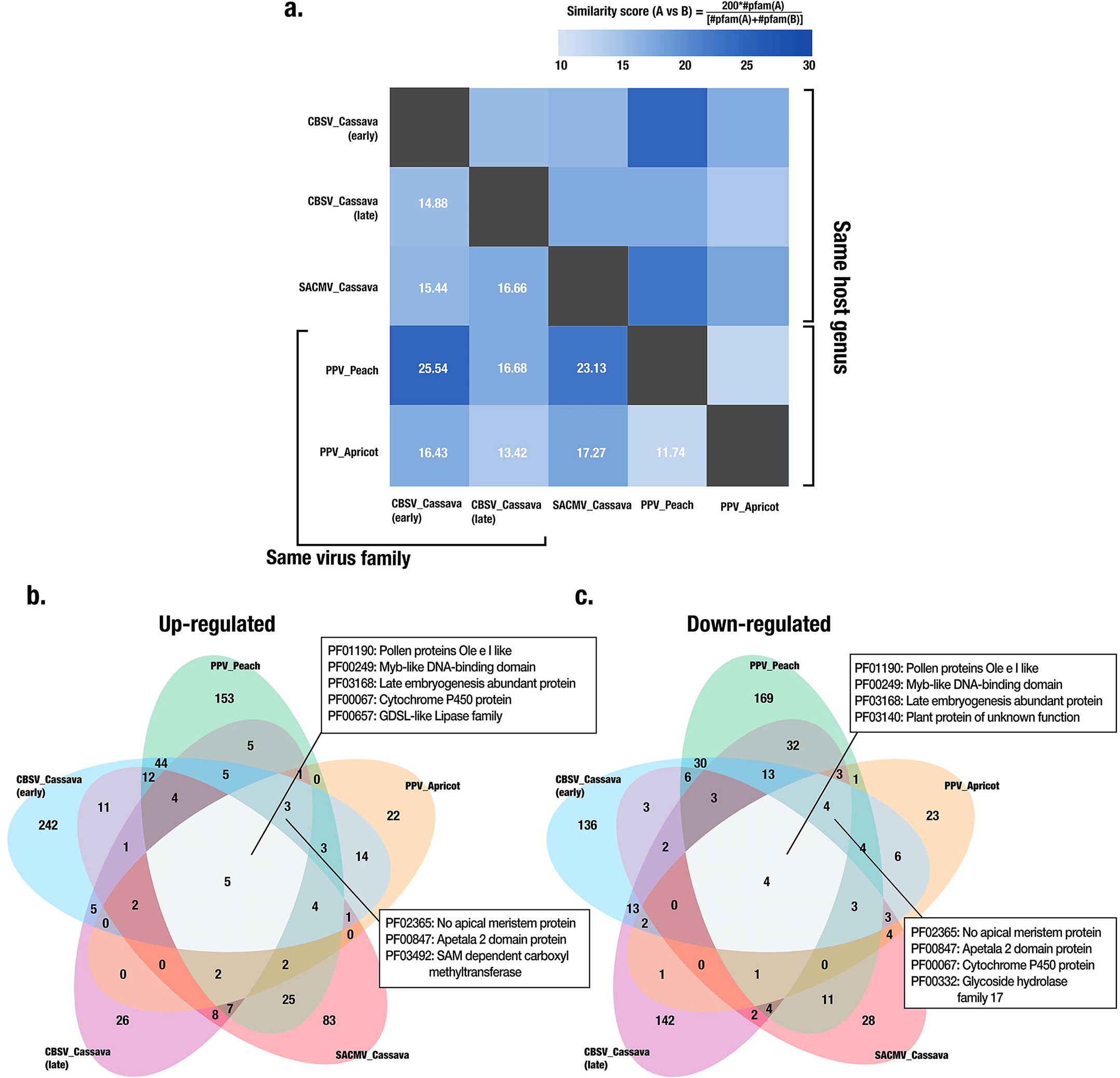
Comparison of transcriptome changes in different virus-host systems. We compared our RNA-Seq of CBSV-infected cassava with three other poty virus-plant transcriptome studies (late-stage cassava-CBSV infection, <link rid="bib34">Maruthi et al., 2014</link>; early-stage peach and late-stage apricot infection with Plum Pox Virus, <link rid="bib49">Rubio et al., 2015a</link>, b). In addition, we included a RNA-Seq study of cassava infected with South African Cassava Mosaic virus, a ssDNA geminivirus pathosystem (<link rid="bib1">Allie et al., 2014</link>), to compare the response of cassava to two different virus types. a. Pairwise similarity matrix of numbers of DEGs in each pathosystem, based on comparison of Pfam identifiers associated with each gene. A pairwise percentage similarity score was computed by multiplying the DEGs in one pathosystem with 200 and then dividing this number by the sum of the DEGs in both pathosystems that were compared. b. and c. Venn diagrams were constructed using the Pfam identifiers associated with DEGs from all five studies. The Fig. shows specific Pfam identifiers common to all five studies as well as those common to all potyvirus studies. b. Up–regulated DEGs. c. Down–regulated DEGs.

We next compared up- and down-regulated DEGs separately across the five datasets (Fig. 5b,c; Table S5) and found five Pfam identifiers in common for up-regulated and four for down-regulated DEGs. Of these, three Pfams that contained genes in both the up- and down-regulated data-sets in all pathosystems included PF01190 (pollen protein Ole e I-like), PF00249 (Myb-like DNA-binding domain) and PF03168 (late embryogenesis-abundant protein). While PF01190 represents a largely uncharacterised protein family, the other two identifiers comprise stress response genes. Genes belonging to PF00067 are up–regulated in all pathosystems and this Pfam comprises the cytochrome P450 protein family, which includes members involved in the synthesis of plant defence compounds. Genes encoding members of the GDSL-like lipase family (PF00657) are also up-regulated in all pathosystems and are required for systemic ethylene-mediated resistance (Kwon et al., 2009; Kim et al., 2013).

In addition to the Pfams common to all RNA-Seq datasets, genes in another three Pfams were found up- and four were down-regulated in all potyvirus infection studies (Fig. 5b,c). Of these, genes belonging to PF02365 (NO APICAL MERISTEM protein) and PF00847 (APETALA 2 domain protein) were found in both the up- and down-regulated data sets.

## DISCUSSION

We show that the transcriptional response of cassava to CBSV infection was significantly stronger in the susceptible 60444 variety than the resistant KBH 2006/18 variety. This is similar to results reported earlier for extreme virus resistance in potato (Goyer et al., 2015). To ensure that our dataset represents the earliest time-point of complete virus infection, we first measured virus titres in the inoculated scions at three time-points until we found consistent infection in all 60444 plants (Fig. S2). This allowed us to analyse changes in gene expression prior to symptom development, and thus early in the infection process. While graft inoculation may delay infection until the graft junction is fully established, we previously demonstrated consistent infection using the top-grafting method (Moreno et al., 2011; Vanderschuren et al., 2012; Anjanappa et al., 2016). Graft virus transmission using field-infected rootstocks or scions was necessary because no infectious CBSV clones are available (Mohammed et al., 2012; Vanderschuren et al., 2012; Wagaba et al., 2013; Anjanappa et al., 2016). Although the lack of detectable virus titres in KBH 2006/18 indicates the absence of viral replication in the resistant cultivar, we observed a transcriptional response to graft-inoculation. It is currently unknown if this is due to the local recognition of viral RNAs and/or proteins, which are mobilized in the vascular tissues after the graft was established, or to an extended systemic response transmitted by the CBSV-susceptible rootstock to the CBSV-resistant scion.

Based on the DEGs in the susceptible variety, 60444, we found that CBSV infection increases the expression of genes encoding SA synthesis enzymes, which likely increases SA levels that activate SA-regulated genes involved in antiviral responses. Our data suggest that in CBSV-infected cassava SA is synthesized via the phenylalanine rather than the isochorismate pathway. Additionally, we found genes encoding proteins of the DCL2/AGO_5_/AGO2-mediated anti-viral silencing response were upregulated in CBSV-infected 60444 but not KBH 2006/18. The silencing response may beamplified as indicated by the strong upregulation of the gene encoding RDR1, but it is also possible that RDR1 assists in the repression of RDR6–mediated silencing amplification (Ying et al., 2010).

The susceptible 60444 and resistant KBH 2006/18 varieties differed strongly in callose deposition at plasmodesmata during CBSV infection. While DEGs associated with virus movement through plasmodesmata were found only in CBSV-infected 60444, callose deposition at plasmodesmata was reduced in 60444 and increased in KBH 2006/18, which could only be partially explained by the detected DEGs. Nevertheless, the results are consistent with the limitation of CBSV movement in KBH2006/18 from stems into leaves (Anjanappa et al., 2016). An efficacious CBSV infection of susceptible 60444 variety thus involves a virus-induced reduction in callose deposition at the plasmodesmata that is absent in the resistant KBH2006/18 variety.

The meta-analysis of our RNA-Seq dataset and other plant-virus RNA-Seq datasets revealed a low level of similarity in host transcriptome responses to different virus types and at different time-points of infection. A previous transcriptome-based meta-analysis of plant-virus interactions compared transcriptional responses in Arabidopsis after infection with different virus species (Rodrigo et al., 2012). Their results indicated a correlation between host transcriptome response and virus phylogeny, with closely related viruses generating transcript regulation of similar host genes. Since our analysis did not involve a model plant, we managed differences in genome annotations in different species by using Pfam identifiers. While Pfam identifiers reduce resolution because they represent protein families instead of specific genes, this approach does not require identification of exact gene orthologs across multiple species. Thus, this allows the comparison of transcriptome regulation between different viruses and plant species. Overall, we found only a small number of common Pfam identifiers, even between different hosts of the same genus or between viruses of the same family. However, based on the numbers of common Pfam identifiers, there was strong temporal impact on gene regulation after virus infection, indicating that earlier transcript-level changes are perhaps most diagnostic to infection by different viruses. Pfam identifiers that were found in common across the selected pathosystems did not relate to classical defence pathways but rather to protein families implicated in developmental processes. A previous review of expression-profile studies also found commonalities within developmental genes across various compatible pathosystems (Whitham et al., 2006). The common regulation of developmental genes might describe similarities in symptom development or downstream signalling processes rather than in primary immune responses across different host-virus interactions.

Together, our study suggests that analysis of early time-points during virus-host interactions captures the emerging impact of virus replication on host gene expression. For example, our RNA-Seq dataset from scion leaves 28 days after grafting revealed a broad immune response (including SA and RNA silencing pathways) in the susceptible cassava 60444 variety that was not found in susceptible cassava plants after a long-term infection (Maruthi et al., 2014). Our meta-analysis of different RNA-Seq datasets also showed significant differences in transcriptome responses at different times after infection. The absence of a detectable immune response in the resistant KBH 2006/18 variety suggests that even earlier time-points after infection should be analysed, preferably using Agrobacterium with infectious CBSV clones or infections of protoplasts followed by transcriptomics or proteomics. The Pfam-based meta-analysis also identified interesting protein families, such as meristem identity proteins (NAM and Apetala2), which were regulated in different host-potyvirus interactions and at different time points after infection. Understanding the functions of these proteins in host–virus interactions might reveal conserved developmental responses to virus infections across different taxa. Our Pfam-based comparison methodology also makes larger scale comparison of crop RNA–Seq studies possible, with the aim of developing a more comprehensive understanding of conserved responses of non–model plants to different biotic and abiotic stresses.

## ACKNOWLEDGEMENTS

The authors would like to thank Asuka Kuwabara for assisting with the microscopy, Kamil Sklodowski for help with the image analysis, and Matthias Hirsch-Hoffman for bioinformatics datamanagement. We also thank Irene Zurkirchen for support in the glasshouse and R. Glen Uhrig for helpful comments on the manuscript. We gratefully acknowledge the support of our research by ETH Zurich. Ravi B. Anjanappa was supported by the Research Fellow Partnership Programme of ETH Global. Devang Mehta is supported by the European Union’s Seventh Framework Programme for research, technological development, and demonstration (EU GA–2013–608422–IDP BRIDGES).

## SUPPORTING INFORMATION

Fig. S1: Experimental scheme for the study. **a.** Schematic representation of the workflow for RNA-seq study. **b.** Workflow for callose quantitation, enzymatic assay and expression analysis for β 1,3 Glucanase.

Fig. S2: CBSV quantitation at 3 different time points after grafting. RT-qPCR quantitation of virus titre (log2 fold change relative to reference gene mePP2A) from individual leaves from 3 independent biological replicates at three different time points after grafting.

Fig. S3: Validation of RNA-seq data by RT-qPCR. RT-qPCR for four genes was performed on the same samples that were used for RNA-seq analysis.

Fig. S4: Virus read counting from unmapped RNA-seq reads. Unmapped reads from each sample sent for RNA-seq were mapped to the two *de novo* assembled virus genomes. **a.** Read distribution across virus genomes for 60444 samples. **b.** Read counts for all samples in the compatible (60444) and incompatible (KBH 2006/18) virus-host interaction.

Fig. S5: RT-qPCR based RDR1 transcript expression across the three time points.

Table S1: Primers used in the study

Table S2: Significantly differentially expressed genes

Table S3: List of denovo assembled transcripts and their differential expression

Table S4: Results of gene set enrichment analysis

Table S5: Comparison of multiple virus-host RNA-seq datasets

Table S6: DEGs in the study that match putative RDR1 targets identified by Cao et al., 2014.

